# Positive selection in the Ebola virus proteome

**DOI:** 10.1101/556910

**Authors:** Pouria Dasmeh

**Affiliations:** Department of Chemistry and Chemical Biology, Harvard University, Cambridge, MA USA 02139.; Department of Biochemistry, Institute for Data Valorization (IVADO), University of Montreal, 2900 Edouard-Montpetit, Montreal, Quebec H3T 1J4, Canada.; Institute for Data Valorization (IVADO), University of Montreal, 2900 Edouard-Montpetit, Montreal, Quebec H3T 1J4, Canada.

## Abstract

To gain insight into the recent evolution of EBOLA virus (EBOV) proteins and identify evolutionary hotspots, I investigated patterns of amino acid substitutions in the proteomes of 49 members of EBOV genus including EBOV, Bundibugyo virus (BDBV), Reston virus (RESTV), Sudan virus (SUDV) and Taï Forest virus (TAFV) spanning outbreaks from 1974 to 2014 in Democratic Republic of Congo (DRC), Uganda, Gabon, Sudan, Ivory Coast, Philippines, and Guinea among humans, non-human primates, and pigs. Of the seven genes of the Ebola genome, I found significant variations in evolutionary rates of Glycoprotein (GP), RNA/dependent RNA polymerase (L), and Nucleoprotein (NP) across different lineages. GP and NP were found to evolve at 5- and 3-fold higher rate than other EBOV genes. Several residues in GP and NP show significant d*N*/d*S* >1 along the internal branches of the phylogenetic tree leading to Philippines and Sudan outbreaks. Most of these residues are located in solvent exposed areas shown previously to be antigenic. We further identified significant changes in specific amino acid properties of GP, NP and L during the recent evolutionary history of EBOV outbreaks and in particular during the 2014 outbreak. The positively selected residues and chemical properties are consistent with changes in epitope interaction and could thus be important in detecting adaptation of EBOV to the host immune defense system.

## Introduction

The recent 2014 Ebola outbreak (EBOV2014) has infected more than 20,000 and claimed the life of more than 8000 people in several African countries [1]. Despite the world-wide efforts to battle the virus, it is predicted that the outbreak might not be fully contained before late 2016 [2]. EBOV has a fatality rate of 50% to 90% due to a combination of misdiagnose [3], poor hygienic maintenance of medical facilities, and in particular the complicated multi-strategy infection mechanism of the virus [4,5]. Whereas the first two problems must be solved by clinical measures, the latter requires molecular insight to counter EBOV adaptation to human hosts.

The negative single stranded RNA (ssRNA) genome of EBOV has seven genes that code for at least ten protein forms including glycoprotein (GP), nucleoprotein (NP), RNA-dependent RNA polymerase (L), matrix protein VP24 and VP40 [6], transcription activator VP30 [7], polymerase co-factor VP35 [8], and a non-structural soluble glycoprotein, sGP [9]. At the core of the virus particles, the EBOV genome is wrapped around viral proteins NP, VP35, VP30 and L, forming a nucleocapsid structure. VP24 and VP40 bridge the nucleocapsid with an outer viral envelope flecked with GP spikes [10], (Figure S1 in File S1). VP24 suppresses interferon production and, along with VP35 and NP, form nucleocapsid structures [11], whereas GP spikes are essential for EBOV pathogenicity, contributing to viral attachment and fusion [12,13].

Early evolutionary analyses demonstrated that EBOV evolve with a rate similar to hepatitis B virus (~3.6×10^−5^ nonsynonymous substitutions per site per year within the GP gene) but hundred times lower than that of retroviruses and human influenza A viruses [14]. Calculations based on whole gene sequences also estimated the rate of evolution to vary from 0.46 × 10^−4^ nucleotide substitutions per site per year for Sudan EBOV to 8.21 × 10^−4^ substitutions per site per year for Reston EBOV [15].

EBOV has previously caused major outbreaks in Sudan, Democratic Republic of Congo (DRC), Republic of Congo and Gabon [16,17]. Therefore, an important concern is the phylogenetic relation between the current 2014 outbreak and the rest of the lineages. The recent EBOV outbreak has been thought to arise from a divergent endemic variant of the Zaire EBOV lineage [18], but the Guinea 2014 lineage was recently placed within the same Zaire EBOV lineage that caused previous outbreaks [19]. EBOV strains from patients in Sierra Leone recently revealed substitution rates roughly twice as high during outbreak as between outbreaks, and mutations were more frequently non-synonymous during outbreak [20]. Thus, outbreaks seem to provide an adaptation opportunity for the virus, and a main concern is whether EBOV evolves under adaptive or neutral evolution.

Despite previous efforts towards understanding evolution of EBOV, it is currently unknown if these changes, on the time scales of outbreaks seen so far, are caused by specific evolutionary host adaptations, as could be feared, or by random evolutionary (i.e. neutral) drift. The purpose of this work is to cast light on this question and identify the proteins, amino acids, and properties that are selected during outbreaks. With the availability of structures of some EBOV proteins, anomalously evolving residues can be structurally mapped providing valuable information for molecular-level targeting of the virus.

To answer these questions, I investigated the patterns of site-resolved evolutionary rate variation across 49 strains of EBOV, Bundibugyo virus (BDBV), Reston virus (RESTV), Sudan virus (SUDV) and Taï Forest virus (TAFV), collected from 1976 to 2014 outbreaks. I find that only three genes, NP, GP, and L, show significant rate variation across different lineages. Interestingly, GP and NP show signatures of positive selection. A branch-by-branch scan of positive selection in the evolutionary history of the virus reveals several residues under positive selection in EBOV lineages of the Philippines and Sudan outbreaks. In both GP and NP, the positively selected residues are positioned in solvent exposed regions shown previously to be antigenic. Finally, our analysis identifies adaptive changes in the chemical properties of GP and NP relevant to future prevention of Ebola adaptation to human hosts.

## Results and discussion

To evaluate the evolutionary rate (ER) of coding sequences of EBOV genes, I used the ratio of the rates of non-synonymous to synonymous substitutions (d*N*/d*S*) [21], which measures the type and strength of selection. The evolution of a protein is potentially under positive selection when the normalized rate of nonsynonymous substitutions (d*N*) significantly exceeds the rate of synonymous substitutions (d*S*), i.e., d*N*/d*S*>1. In contrast, d*N*/d*S* <1 implies purifying selection with non-synonymous substitutions less common than expected, compared to synonymous substitutions. When proteins evolve by random drift (neutrally), d*N*/d*S* is expected to be ~ 1.

However, whole-gene estimation of d*N*/d*S* might not fully capture the selection pressure because of variations in ER at different residues as well as branches of the phylogenetic tree. For example, positive selection (i.e., d*N*/d*S*>1) in specific residues could be masked by the presence of highly conserved (i.e., d*N*/d*S*<<1) or neutrally evolving (i.e., d*N*/d*S*<1) regions in proteins. To remove these biases, I employed different types of evolutionary analyses as explained in Methods.

I first estimated pairwise d*N*/d*S* for EBOV proteins. Since d*N*/d*S* estimation is highly sensitive to saturation of synonymous substitutions (i.e., d*S* > 1), I only considered comparisons in which d*S* < 1. As shown in Figure 1A and presented in File S1, Table S1, the median of pairwise d*N*/d*S* are 0.25, 0.15, 0.06, 0.04, 0.05, 0.07 and 0.05 for GP, NP, L, VP24, VP30, VP35 and VP40, respectively. Thus, I find that GP and NP on average have had ~5- and ~3-fold higher rates of evolution compared to the remaining EBOV proteins.

**Figure 1.**
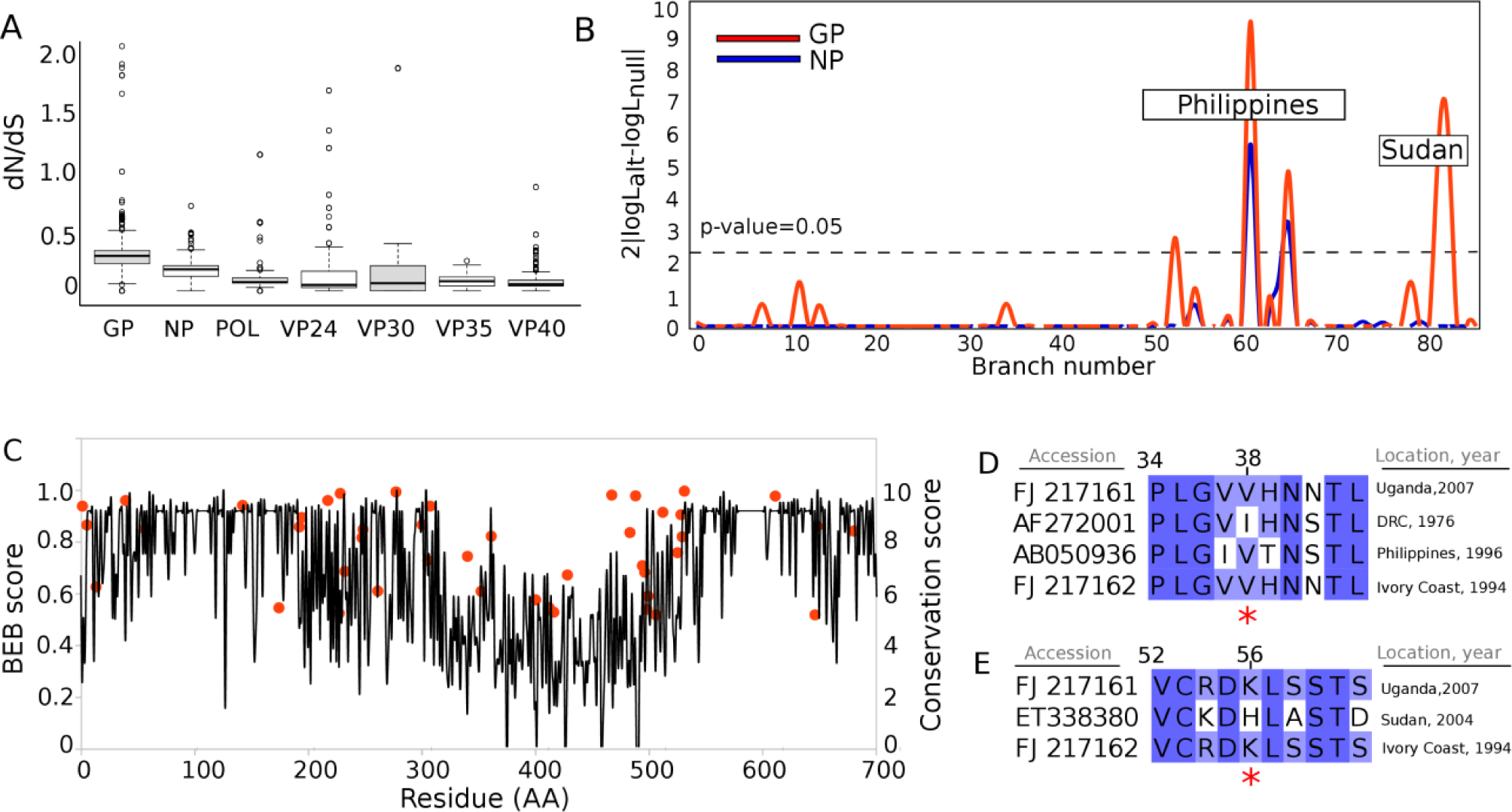
Evolutionary rate is higher in Glycoprotein (GP) and Nucleoprotein (NP) than other EBOV proteins. A) Pairwise estimated d*N*/d*S* for EBOV proteins. Comparisons with dS>1 were removed to avoid synonymous saturation bias. Center lines show the medians; box limits indicate the 25th and 75th percentiles; whiskers extend 1.5 times the interquartile range from the 25th and 75th percentiles, and outliers are represented by dots. n = 385, 358, 364, 347, 341, 333, 352 sample points for GP, NP, L, VP24, VP30, VP35 and VP40 respectively. B) Selection criteria, twice the difference in logarithm of likelihood between alternative and null models of branch-site positive selection, versus branch number for GP (red) and NP (blue). Dashed line shows 0.05 significance level. C) Bayes Empirical Bayes score for positively selected residues (red circles) and conservation score for all residues (gray line) versus residue number in GP. Multiple sequence alignment of D) residues 34−43 and E) residues 52−61 in GP are shown with positively selected residues marked with red asterisks.

Pairwise d*N*/d*S* may not reveal true selection on coding sequences because of averaging effects. In particular, although genes evolving with d*N*/d*S*<1 are likely under purifying selection leading to residue conservation, it is known from both real biological cases and simulations that positive selection can occur in specific sites while whole-gene d*N*/d*S* is as low as 0.25 [22,23,24]. I thus studied the evolution of the EBOV proteome using models that account for d*N*/d*S* variation along the branches of EBOV phylogeny or across sequences of the proteins, to identifyevolutionary hotspots within the EBOV proteome.

To systematically investigate patterns of d*N*/d*S* variation in EBOV proteins, I first checked whether whole-gene d*N*/d*S* is significantly different among branches of the EBOV phylogeny (Figure S2 in File S1), i.e., whether the virus have evolved with different rates during different outbreaks. I compared the likelihood of one-ratio (M0) versus free-ratio (FR) models (File S1, Table S2). LRT is only significant for NP, GP and L genes in this comparison with p-values of 2.15×10^−7^, 2.58×10^−10^, and < 10^−15^, respectively. I then asked whether d*N*/d*S* is variable across the sequence and the functional domains of EBOV proteins are under positive selection throughout the whole phylogeny (see the explanation of M7/M8 and M8/M8fix model comparisons in File S1). From Table S3 in File S1, GP and L proteins show significant d*N*/d*S*>1 in certain residues with p-values of 2.4×10^−3^ and 5.5×10^−4^, respectively (see Table S4 in File S1). However, only L protein passes the strictest criteria for positive selection (p ~ 5.5×10^−4^, see M8/M8fix model comparison in File S1 and Table S3 in File S1).

Although even the most strict site-model test only found positive selection in the L protein, real evolutionary rates more commonly change locally and rapidly for specific sites under positive selection or mutation-selection balance [25]. To identify such more realistic site-specific selection, I performed branch-site test to detect positive selection for each branch of the phylogenetic tree with no assumption of equal evolutionary rates and d*N*/d*S* allowed to vary along the branch of interest and across the sequence, as will be the case in real proteins. The likelihood of this model was compared with the null model with d*N*/d*S* = 1 and a p-value was calculated using the relevant likelihood ratio test [26]. In addition, one can compare likelihood of the null of model of branch-site test with that of M1 model. Since M1 model assumes strict neutrality, a significant likelihood difference in this comparison shows the excess of d*N*/d*S*~1 along the branch of interest which is interpreted as the sign for relaxation of functional constraints [27]. I would thus be able to distinguish between natural and relaxed evolution of EBOV proteins using this approach.

As shown in Tables S5 and S6 in File S1 and S8 in File S2, branch-site tests for positive selection revealed several residues to be under positive selection during the evolution of GP and NP (Figure 1B). GP and NP have both been subject to positive selection in the lineages of the 1990, 1996, 2008 and 2009 EBOV outbreaks in the Philippines. In addition, GP underwent positive selection during the 1976, 1979, and 2004 outbreaks in Sudan, rendering this protein subject to most branch-specific positive selection. Simply put, GP accepts non-synonymous substitutions more frequently than other EBOV proteins, compared to the synonymous background.

I presented all residues under positive selection in GP and NP proteins in File S1, Table S6. From the table, two interesting features become apparent: First, there is no overlap between residues detected by M7/M8 comparison (File S1, Table S4) and branch-site test for positive selection (File S1, Table S6). Therefore, GP is under weaker positive selection throughout the entire phylogeny than within specific lineages. Second, I observe the occurrence of positive selection in adjacent residues such as 192 and 194, 227 and 228, 247 and 248, 305 and 307,494 and 496, 498 and 499, 528 and 529 and, 646 and 648. This clearly shows that positive selection most likely occurs in important regions of the protein where multiple substitutions are necessary e.g. in functional adaptation, similar to previous observations [28,29].

It is important to rule out the erroneous detection of positively selected residues because of poor alignments. For example, false detection of positively selected residues can amount to 50−55% in poorly aligned regions [30]. Therefore, I plotted residues with BEB scores > 0.5 along with residue conservation scores for the 49 EBOV GP sequences (see Methods for detail) (Figure 2A). Sequence conservation is relatively high except for residues 312−513 where a highly variable region is observed. Positively selected residues with lower BEB scores (>0.5 and <0.8) are also enriched in this region. Although this region of high variability is robust to alignment, I report positively selected residues within this region with caution. Nonetheless, residues with BEB scores > 0.8 are all embedded in highly conserved regions (e.g., 39, 56, 142 and 217 with BEB scores of 0.96, 0.85, 0.94 and, 0.96 as shown in Figures 2D-E and Figure S3 in File S1).

**Figure 2.**
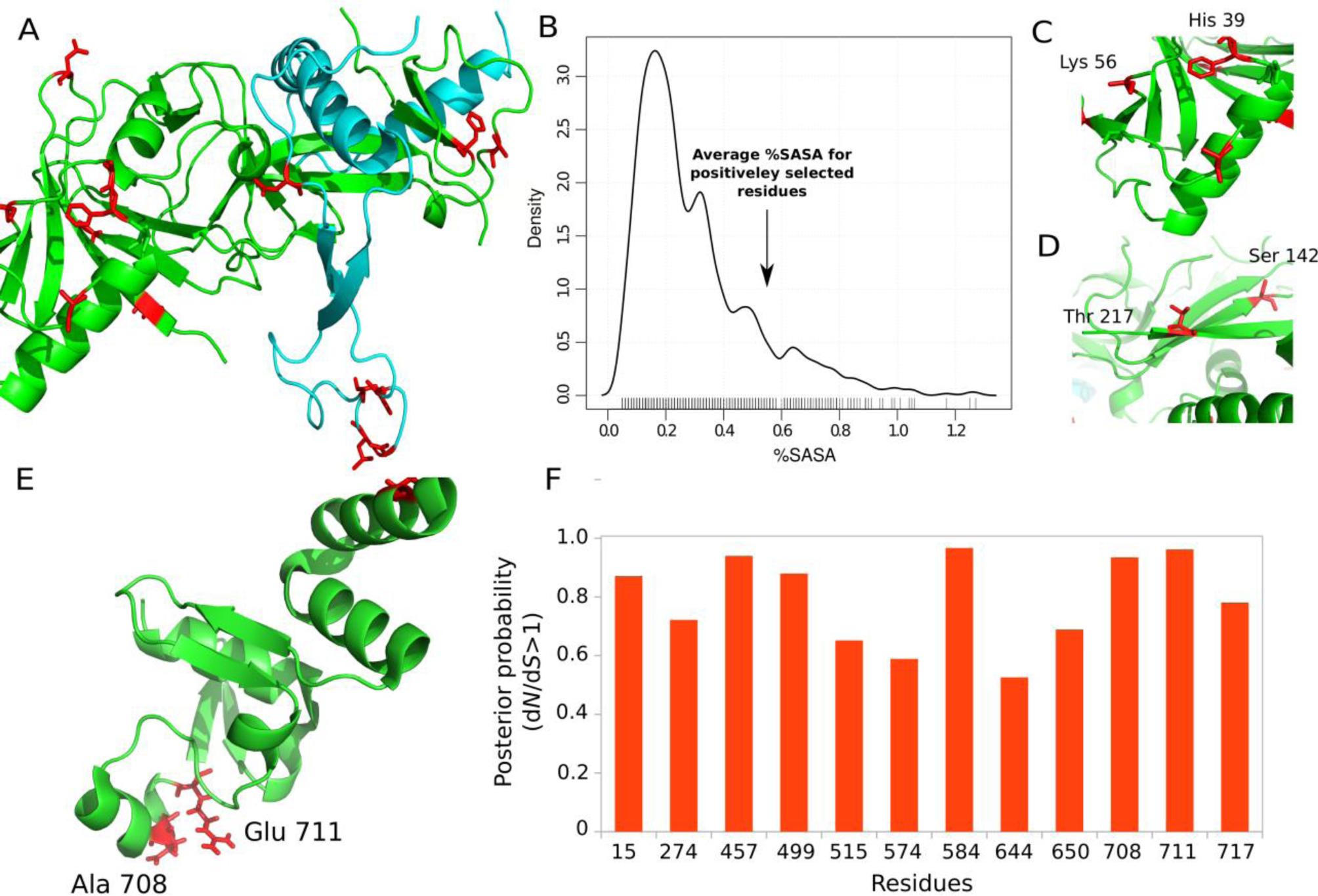
Positively selected residues in EBOV Glycoprotein (GP) and Nuleoprotein (NP) are located in solvent exposed regions. A) Crystal structure of GP1 (in green) and GP2 (in cyan) subunits of EBOV GP (PDB ID= 3CSY) with positively selected residues shown in red. B) Density plot of percentage of solvent accessible surface area (%SASA) of residues within the protein to fully random coil configuration (i.e., SASA_protein_/SASA_coil_). Different residues such as C) His39 and Lys56 and D) Ser142 and Thr217 could potentially lie within conformational epitopes. E) Crystal structure of the C-terminal of EBOV NP (PDB ID= 4QB0) with two positively selected residues Ala708 and Glu711 shown in red. F) BEB scores of different codon positions throughout the NP sequence.

I also investigated whether positive selection in GP and NP has occurred in internal branches of the phylogenetic tree, i.e. representing fixed mutations, or simply in mutations that could be purged from population by later evolution of the virus. To this end, I resolved substitutions on each branch of the phylogenetic tree for GP and NP proteins using ancestral reconstruction approach based on maximum likelihood estimations (see Methods). The most probable ancestral amino acids in GP and NP are shown on each node of the phylogenetic tree for GP and NP in Figure sets S4 and S5 in File S3. From the figures, almost all positively selected residues are fixated along internal branches or in different clades of the phylogenetic tree. For example, an Asn to Glu mutation at position 227 in GP is positively selected in 2008 and 2009 RESTV outbreaks in Philippines. Another example is positive selection of Thr30 in the branch leading to EBOV strains in Philippines, Uganda and Sudan.

The branch-site test for positive selection explicitly detects residues with d*N*/d*S* >1 along different branches of the phylogenetic tree. Therefore, the observation of positive selection is independent of elevated d*N*/d*S* due to enhancement of substitutions with d*N*/d*S*~1 often interpreted as the sign for relaxation of functional constraints. I therefore examined the extent to which EBOV evolves under this distinct mode of evolution (Table S8 in File S2). Surprisingly, all EBOV proteins show relaxed evolution along different branches of the phylogenetic tree. Proteins L and GP have the maximum number of branches under relaxed evolution (i.e., with 16 and 13 branches respectively). Among the outbreaks, the Philippines had the maximum number of proteins under relaxed evolution (i.e., all proteins except VP24). Significant relaxation is also observed within the outbreak of Sudan.

One possible reason for the special evolution of GP in different lineages might be the role of this protein in the first and most important step in the infection mechanism of EBOV. GP is in fact involved in attachment and membrane fusion of EBOV [12,13]. Moreover, it has been shown that multiple immunoglobin (IgG) antibodies bind to GP epitopes, which permits binding of complement component 1R (C1) to the fragment crystalized (Fc) regions of antibodies and hence contribute to antibody-dependent infection [4]. This complex stabilizes the interaction between EBOV and its receptor, therefore enhancing the infection4.

Figure 2A-B show the residues under positive selection mapped onto the crystal structure of EBOV GP and the average value of solvent accessible surface area (SASA) of these residues compared with other residues in a density plot. The majority of positively selected residues are located on the surface of GP, on average ~50% exposed to solvent and other proteins. Surprisingly, positively selected residues 217, 228, 300, 305, 307, 340, 395, and 409 with high BEB scores (Figure 2A) are exactly within the three regions in the protein, i.e., residues 216−239, 301−359, and 381−411, that were previously shown to be immunogenic regions of EBOV GP recognized by anti-EBOV IgG [31]. Among these, residues 217, 228, 300, 305 and 307 are within highly reliable sequence alignments and thus prime candidates for positive selection in GP epitopes. I also find positively selected residues that are adjacent in the 3D structure (residues 39 and 56 in Figure 2C and residues 142 and 217 in Figure 2D) possibly indicating coevolution (epistasis) of conformational epitopes of GP. Figure 2E shows the crystal structure of the C-terminal domain of EBOV NP with positively selected residues mapped onto the structure. Similar to GP, residues with high BEB scores shown in Figure 2F (e.g. residues 708 and 711) tend to occur in solvent accessible regions of the protein. I also observe that positively selected residues 499 and 515 are located in previously reported reactive epitopes of NP [31].

I previously argued that the detected positive selection could be due to selection for biophysical properties or specific structural features in proteins [23,24], thus, I then determined if the positively selected sites in EBOV are related to specific protein properties. To this aim, I analyzed whether amino acid properties have changed significantly during the recent evolution of the EBOV proteome, using the TreeSAAP method [47] (see File S1 for details of calculations). In brief, TreeSAAP evaluates significant changes in amino acid properties during protein evolution. Since d*N*/d*S* estimation is ultimately a counting approach and does not distinguish between different types of non-synonymous substitutions, TreeSAAP complements the d*N*/d*S* analysis.

As summarized in Table S7 in File S1, GP, NP and L proteins again show significant selection (*p* < 5×10^−2^ and 10^−3^) in specific amino acid properties throughout the phylogenetic history. In particular, GP has been subject to major changes in chemical properties that are statistically significant compared to random evolution. Electrostatic properties such as polar requirements and isoelectric point, but also alpha helix propensity and hydrophobicity have changed significantly (Table S7 in File S1). These properties could relate to the function and stability of GP during host membrane interaction. NP and L also had notable properties such as polarity, hydropathy and alpha-helical tendencies changing significantly during the recent evolution of EBOV. Table S9-S11 in File S2, show codons detected by TreeSAAP to be under positive-destabilizing selection in GP, NP and L proteins in different branches of the phylogeny along with properties with significant changes (p-value < 0.001). In GP, 20 out of 44 positively selected residues (i.e., dN/dS>1) show significant changes in amino acid properties. In NP and L, 8 out of 12 and all of the positively selected residues are detected by TreeSAAP, validating our identification of evolutionary hotspots in EBOV proteins

By scanning the sequences of GP, NP, and L by sliding-windows, the substitutions affecting these properties seem to fall in non-overlapping regions. Figure 3A shows such a sequence-property plot for the alpha helix propensity and isoelectric point of different codons within the EBOV GP sequence. I also showed the 3D location of two peaks of this plot in the structure of EBOV GP (named as α and β clusters) in Figure 3B with sequences shown in Figures 3C and 3D, respectively. Substitutions such as Q28H, T30V, T30A and S32P increase the isoelectric point of the protein. Surprisingly, alpha helix propensity peaks at residues 75-81 (i.e., the β cluster) where a short helix is located in EBOV GP structure. Alpha helix propensity is strongly increased by substitutions such as S76A and P80L. The accelerated evolution of three proteins GP, NP and L within EBOV that I have identified may therefore relate to changes in surface interaction propensities, as most importantly seen in the case of GP, as well as structural stability of membrane fusions required for proper function of these proteins.

**Figure 3.**
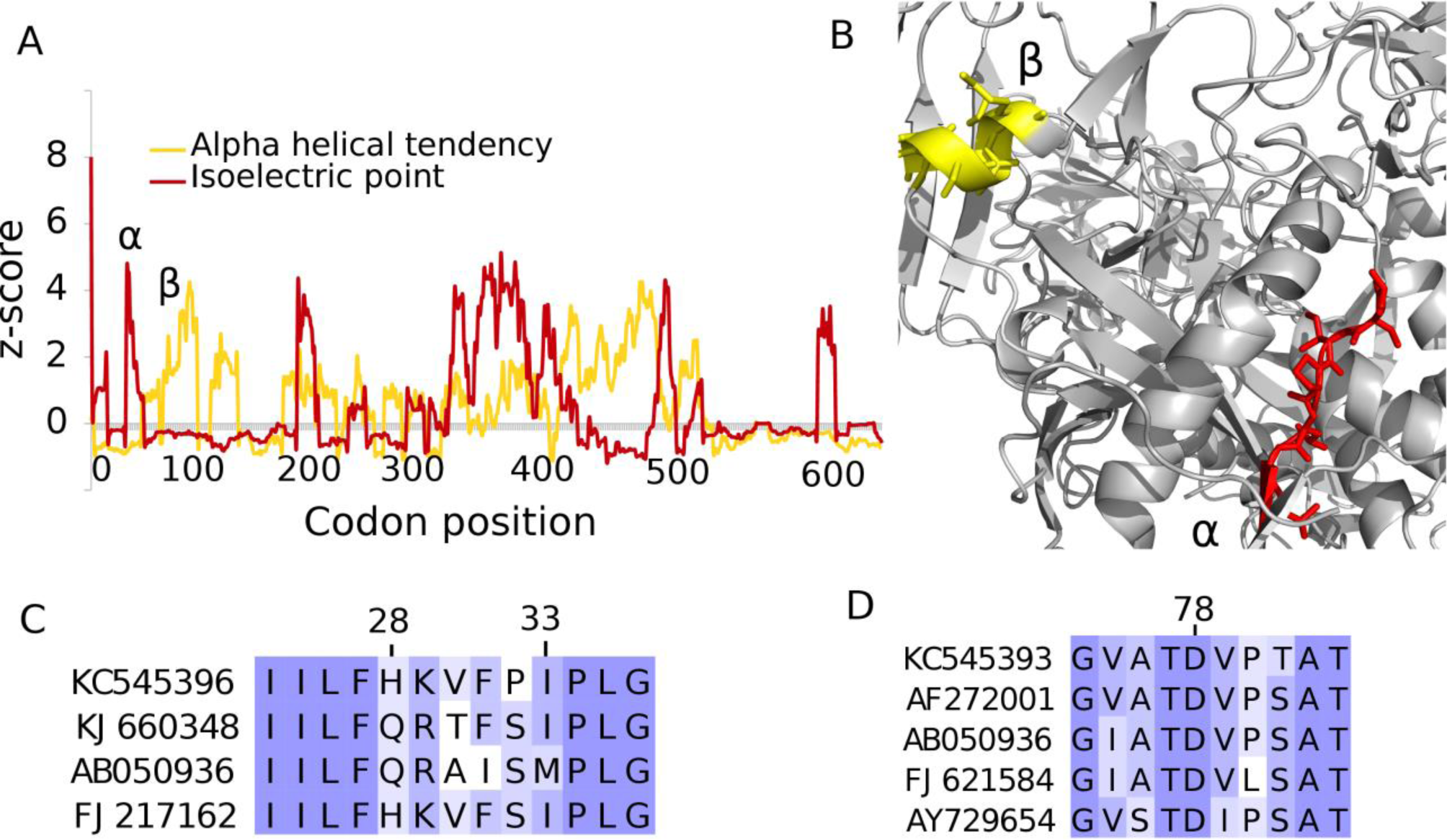
Ebola virus Glycoprotein shows signatures of positive selection in amino acid properties relevant for function as well as folding stability. A) Z-score for sliding window analysis of alpha helical tendency (shown in yellow) and isoelectric point (shown in red) throughout EBOV GP sequence. Z-scores greater than 3.09 correspond to p-values < 0.001. Window size is set to 15 codons. B) α and β clusters of residues from panel A mapped onto the crystal structure of GP (PDB ID =3CSY). C) The α cluster consists of residues 28 to 33 which shows significant changes in isoelectric point. D) In β cluster, from residues 75-81, substitutions enhance alpha helical tendency.

Using the d*N*/d*S* analysis, we did not observe significant positive selection in the lineage leading to 2014 EBOV species (i.e, KJ660346-348 strains in the dataset) as speculated by Gire et al [20]. However, despite the insignificant likelihood ratio test for detection of positive selection, I observed that residue 291 has a BEB score of 0.98 consistent with a Trp to Arg mutation in the Gueckedou-C05 strain of 2014 Guinea outbreak [32]. Although such a charge-changing substitution could potentially induce functional changes, the absence of positive selection might be due to the fact that the time scale of this latest outbreak is still too short for any major adaptations to have become fixated. Since the evolutionary rate is higher during outbreaks, prolonged outbreaks are likely to enable adaptation opportunities towards human hosts.

Our finding of positive selection of GP during the recent evolution of EBOV is in line with the recent finding that GP is the only protein that has accumulated only nonsynonymous substitutions within EBOV genomes of Sierra Leone patients^20^. In addition, Martinez et al. found a GP mutant, F88A, to facilitate entry into various human cell types [33]. Furthermore, Azizian et al. reported positive selection in the site 429 of GP [34] (corresponding to the site 431 in our sequence alignment; see File S1) also detected using M7/M8 comparison in our approach. Together with our analysis, these results establish that GP is the prime candidate for adaptation of EBOV, and the surface-centered GP properties and amino acids that I identify as adaptively changed may help in attempts to halt future adaptation of the virus to human hosts.

## Methods

I retrieved coding sequences of 49 EBOV genomes with the inferred Bayesian phylogenetic tree from the recent study of Dudas and Rambaut [19] (Figure S2 in File S1). These sequences constitute a sampled dataset of genome sequences from the genus *Ebolavirus* spanning major previous outbreaks in Democratic Republic of Congo (DRC), Uganda, Gabon, Sudan, Ivory Coast, Philippines and Guinea. One could include more viral genomes from patients infected during the 2014 outbreak. However, selection analysis, and in particular d*N*/d*S* estimation, is only accurate when the lineages are sampled in balance to avoid bias, [35] and the 49 genomes used here are thus those relevant for our purpose.

The original concatenated sequences were divided into seven protein alignment files. Protein alignments were further processed with the codon-based CLUSTAL algorithm [36] implemented in MEGA [37]. I used the ratio of the rates of nonsynonymous to synonymous substitutions (i.e., d*N*/d*S*) as a measure of selection strength as commonly applied [38]. To calculate the maximum likelihood (ML)-based d*N*/d*S,* I employed the CODEML program within the PAML suite [39]. The equilibrium codon frequencies were estimated from the products of the average observed frequencies in the three-codon positions (F3X4 model). For each of the seven EBOV genes, the tree branch lengths were first estimated with the M0 model that assumes one d*N*/d*S* across all branches and the branch length estimates were used for more advanced codon models. Ancestral sequences were inferred using the Maximum Likelihood method [40] implemented in MEGA5 [41] under the JTT model [42] as the model with highest Bayesian Information Criterion (BIC) score [43].

Different types of d<u>*N*</u>/d*S* analysis were performed. First, I calculated the whole-gene d*N*/d*S* between protein sequences of any two species of interest using pairwise analysis. Second, branch models estimatd d*N*/d*S* along branches of EBOV phylogenetic tree while assuming an average ER for different residues. Third, a set of site models (i.e., M1, M2, M3, M7, M8 and M8fix models, as explained in File S1) were used to estimate d*N*/d*S* variation at different residues while taking an averaged ER of the virus along each branch of the phylogenetic tree. Eventually, a branch-site model test was employed to evaluate d*N*/d*S* for few amino acid residues along specific branches of interest in the phylogenetic tree of the virus using in house shell scripts. I employed maximum likelihood (ML) estimation of d*N*/d*S* via codon models (Material and methods). For all tested hypotheses, I compared the likelihoods of null and alternative models with likelihood ratio tests [44]. All the d*N*/d*S* estimations were performed using the CODEML package within the PAML suite [45]. Models were compared in terms of statistical significance using likelihood ratio tests [46] and posterior probabilities of residues under positive selection were identified using the Bayes Empirical Bayes (BEB) method for branch site tests of positive selection.

Selection analyses on amino acid properties were performed using the TreeSAAP approach [47]. For GP, I computed the solvent accessible surface area (SASA) from the crystal structure (PDB ID= 3CSY) [48] using the POPS algorithm [49]. Alignment conservation scores were calculated by the AMAS method [50] implemented in JALVIEW [51] which measures number of physiochemical properties conserved in each column of alignment. All codon numbers refer to alignment positions. For a correspondence between the alignment positions referred throughout this study and the real sequences see File S1. Statistics was performed using the R package [52] and BoxPlotR web tool [53].

## Acknowledgment

The computations in this paper were run on the Odyssey cluster supported by the FAS Division of Science, Research Computing Group at Harvard University.

## References

1 Team WER (2014) Ebola virus disease in West Africa—the first 9 months of the epidemic and forward projections. N. Engl. J. Med. 371: 1481–95.

2 Lewnard JA, Mbah MLN, Alfaro-Murillo JA, Altice FL, Bawo L, Nyenswah TG, Galvani AP (2014) Dynamics and control of Ebola virus transmission in Montserrado, Liberia: a mathematical modelling analysis. Lancet Infection Dis. 14: 1189–1195.

3 Upadhyay DK, Sittig DF, Singh H (2014) Ebola US Patient Zero: lessons on misdiagnosis and effective use of electronic health records. Diagnosis. DOI: 10.1515/dx-2014-0064.

4 Takada A, Feldmann H, Ksiazek TG, Kawaoka Y (2003) Antibody-dependent enhancement of Ebola virus infection. J. Vir. 77: 7539–7544.

5 Misasi J, Sullivan NJ (2014) Camouflage and misdirection: the full-on assault of Ebola virus disease. Cell 159: 477–486.

6 Timmins J, Schoehn G, Ricard-Blum S, Scianimanico S, Vernet T, Ruigrok RW, Weissenhorn W (2003) Ebola virus matrix protein VP40 interaction with human cellular factors Tsg101 and Nedd4. J. Mol. Biol. 326: 493–502.

7 Modrof J, Becker S, Muhlberger E (2003) Ebola virus transcription activator VP30 is a zinc-binding protein. J. Vir. 77: 3334–3338.

8 Prins KC, Cárdenas WB, Basler CF (2009) Ebola virus protein VP35 impairs the function of interferon regulatory factor-activating kinases IKKε and TBK-1. J. Vir. 83: 3069–3077.

9 Simmons G, Wool-Lewis RJ, Baribaud F, Netter RC, Bates P (2002) Ebola virus glycoproteins induce global surface protein down-modulation and loss of cell adherence. J. Vir. 76: 2518–2528.

10 Lee JE, Fusco ML, Hessell AJ, Oswald WB, Burton DR, Saphire EO (2008) Structure of the Ebola virus glycoprotein bound to an antibody from a human survivor. Nature 454: 177–182.

11 Hoenen T, Groseth A, Kolesnikova L, Theriault S, Ebihara H, Hartlieb B et al. (2006) Infection of naive target cells with virus-like particles: implications for the function of ebola virus VP24. J. Vir. 80: 7260–7264.

12 Yonezawa A, Cavrois M, Greene WC (2005) Studies of ebola virus glycoprotein-mediated entry and fusion by using pseudotyped human immunodeficiency virus type 1 virions: involvement of cytoskeletal proteins and enhancement by tumor necrosis factor alpha. J. Vir. 79: 918–926.

13 Volchkov VE, Feldmann H, Volchkova VA, Klenk HD (1998) Processing of the Ebola virus glycoprotein by the proprotein convertase furin. Proc. Natl. Acad. Sci. U.S.A. 95: 5762–5767.

14 Suzuki Y, Gojobori T (1997) The origin and evolution of Ebola and Marburg viruses. Mol. Biol. Evol. 14: 800–806.

15 Carroll SA, Towner JS, Sealy TK, McMullan LK, Khristova ML, Burt FJ et al. (2013) Molecular evolution of viruses of the family Filoviridae based on 97 whole-genome sequences. J. Vir. 87: 2608–2616.

16 Georges-Courbot MC, Sanchez A, Lu CY, Baize S, Leroy E, Lansout-Soukate J et al. (1997) Isolation and phylogenetic characterization of Ebola viruses causing different outbreaks in Gabon. Emerg. Infect. Dis. 3: 59.

17 Leroy EM, Baize S, Mavoungou E, Apetrei C (2002) Sequence analysis of the GP, NP, VP40 and VP24 genes of Ebola virus isolated from deceased, surviving and asymptomatically infected individuals during the 1996 outbreak in Gabon: comparative studies and phylogenetic characterization. J. Gen. Virol. 83: 67–73.

18 Baize S, Pannetier D, Oestereich L, Rieger T, Koivogui L, Magassouba NF et al. (2014) Emergence of Zaire Ebola virus disease in Guinea. N. Engl. J. Med, 371: 1418–1425.

19 Dudas G, Rambaut A (2014) Phylogenetic analysis of Guinea 2014 EBOV Ebolavirus outbreak. PLoS currents 6.

20 Gire SK, Goba A, Andersen KG, Sealfon RS, Park DJ, Kanneh L et al. (2014) Genomic surveillance elucidates Ebola virus origin and transmission during the 2014 outbreak. Science 345: 1369–1372.

21 Yang Z, Bielawski JP (2000) Statistical methods for detecting molecular adaptation. Trends Ecol. Evol. 15: 496–503.

22 Swanson WJ, Wong A, Wolfner MF, Aquadro CF (2004) Evolutionary expressed sequence tag analysis of Drosophila female reproductive tracts identifies genes subjected to positive selection. Genetics 168: 1457–1465.

23 Dasmeh P, Serohijos AW, Kepp KP, Shakhnovich EI (2013) Positively selected sites in cetacean myoglobins contribute to protein stability. PLoS Comp. Biol. 9: e1002929.

24 Dasmeh P, Serohijos AW, Kepp KP, Shakhnovich EI (2014) The influence of selection for protein stability on dN/dS estimations. Genome Bio. Evol. 6: 2956–2967.

25 Yang Z, Nielsen R (2002) Codon-substitution models for detecting molecular adaptation at individual sites along specific lineages. Mol. Bio. Evol. 19: 908–917.

26 Yang Z, Dos Reis M (2011) Statistical properties of the branch-site test of positive selection. Mol. Biol. Evol. 28: 1217–1228.

27 Zhang J, Nielsen R, Yang Z (2005) Evaluation of an improved branch-site likelihood method for detecting positive selection at the molecular level. Mol. Biol. Evol. 22.12: 2472–2479.

28 Kosiol C, Vinař T, da Fonseca RR, Hubisz MJ, Bustamante CD, Nielsen R, Siepel A. (2008) Patterns of positive selection in six mammalian genomes. PLoS genetics 4: e1000144.

29 Eikenboom JC, Vink T, Briet E, Sixma JJ, Reitsma PH (1994) Multiple substitutions in the von Willebrand factor gene that mimic the pseudogene sequence. Proc. Natl. Acad. Sci. U.S.A 91: 2221–2224.

30 Markova-Raina P, Petrov D (2011) High sensitivity to aligner and high rate of false positives in the estimates of positive selection in the 12 Drosophila genomes. Genome res. 21: 863–874.

31 Becquart P, Mahlakõiv T, Nkoghe D, Leroy EM (2014) Identification of continuous human B-cell epitopes in the VP35, VP40, nucleoprotein and glycoprotein of Ebola virus. PloS one 9, e96360.

32 Baize S, Pannetier D, Oestereich L, Rieger T, Koivogui L, Magassouba NF et al. (2014) Emergence of Zaire Ebola virus disease in Guinea. New Eng. J. Med, 371: 1418–1425.

33 Martinez O, Ndungo E, Tantral L, Miller EH, Leung LW, Chandran K, Basler CFA (2013) mutation in the Ebola virus envelope glycoprotein restricts viral entry in a host species-and cell-type-specific manner. J. Vir. 87: 3324–3334.

34 Azarian, Taj, et al. (2015) Impact of spatial dispersion, evolution, and selection on Ebola Zaire Virus epidemic waves. Sci. Rep. 5.

35 Kryazhimskiy S, Plotkin J (2008) The population genetics of dN/dS. PLoS genetics 4.12: e1000304.

36 Thomopson J, Higgins DG, Gibson T (1994) Clustal W. Nuc. Acids Res. 22: 4673–4680.

37 Tamura K, Peterson D, Peterson N, Stecher G, Nei M, Kumar S (2011) MEGA5: molecular evolutionary genetics analysis using maximum likelihood, evolutionary distance, and maximum parsimony methods. Mol. Biol. Evol. 28: 2731–2739.

38 Kimura M (1977) Preponderance of synonymous changes as evidence for the neutral theory of molecular evolution. Nature 267: 275–276.

39 Yang Z (2007) PAML 4: phylogenetic analysis by maximum likelihood. Mol. Biol. Evol. 24: 1586–1591.

40 Nei M, Kumar S (2000) Molecular Evolution and Phylogenetics. Oxford University Press, New York.

41 Tamura K et al. (2011) MEGA5: molecular evolutionary genetics analysis using maximum likelihood, evolutionary distance, and maximum parsimony methods. Mol. Biol. Evol. 28.10: 2731–2739.

42 Jones DT, Taylor WR, Thornton JM (1992) The rapid generation of mutation data matrices from protein sequences. Computer applications in the biosciences: CABIOS, 8(3): 275–282.

43 Schwarz G (1978) Estimating the dimension of a model. The annals of statistics 6.2 (1978): 461–464.

44 Yang Z (1998) Likelihood ratio tests for detecting positive selection and application to primate lysozyme evolution. Mol. Biol. Evol. 15: 568–573.

45 Yang Z (1997) PAML: a program package for phylogenetic analysis by maximum likelihood. CABIOS 13: 555–556.

46 Nielsen R, Yang Z (1998) Likelihood models for detecting positively selected amino acid sites and applications to the HIV-1 envelope gene. Genetics 148: 929–936.

47 Woolley S, Johnson J, Smith MJ, Crandall KA, McClellan DA (2003) TreeSAAP: selection on amino acid properties using phylogenetic trees. Bioinformatics 19: 671–672.

48 Lee JE, Fusco ML, Hessell AJ, Oswald WB, Burton DR, Saphire EO (2008) Structure of the Ebola virus glycoprotein bound to an antibody from a human survivor. Nature 454: 177–182.

49 Cavallo L, Kleinjung J, Fraternali F (2003) POPS: a fast algorithm for solvent accessible surface areas at atomic and residue level. Nuc. Acids Res. 31: 3364–3366.

50 Livingstone CD, Barton GJ (1993) Protein sequence alignments: a strategy for the hierarchical analysis of residue conservation. CABIOS 9: 745–756.

51 Clamp M, Cuff J, Searle SM, Barton GJ (2004) The jalview java alignment editor. Bioinformatics 20: 426–427.

52 Statistical Package, R. (2009) R: A language and environment for statistical computing.” Vienna, Austria: R Foundation for Statistical Computing.

53 Spitzer M, Wildenhain J, Rappsilber J, Tyers M (2014) BoxPlotR: a web tool for generation of box plots. Nature methods 11: 121–122.

